# 2′-Deoxyuridine-promoted infection in *Pyricularia oryzae* is counteracted by bacterial thymidine phosphorylase

**DOI:** 10.64898/2026.04.09.717465

**Authors:** Haowei Hu, Hiroka Maeshima, Rikuto Tsukahara, Akira Ashida, Keiko Kano, Emi Mishiro-Sato, Sotaro Chiba, Daigo Takemoto, Ikuo Sato

**Author notes:** Co-Corresponding Authors: Ikuo Sato and Daigo Takemoto.

## Abstract

Successful infection by the rice blast fungus *Pyricularia oryzae* depends on precise developmental transitions on the plant surface, yet the extracellular metabolites that regulate these events remain poorly understood. One such metabolite, 2′-deoxyuridine (dU), has been identified as a self-produced infection-promoting factor, but its mode of action has remained unclear. Exogenous dU did not significantly affect conidial germination but accelerated early appressorium initiation and promoted appressorium maturation, as indicated by increased glycogen mobilization and elevated intracellular turgor. Extracellular dU was detected during early infection-related development, indicating that dU accumulates under conditions conducive to appressorium formation. To test whether microbial turnover of dU influences pathogenicity, dU-degrading bacteria were isolated from rice field environments, and *Enterobacter* sp. strain C3 was identified as the most active isolate. Biochemical and structural analyses identified the responsible enzyme as the thymidine phosphorylase DeoA, which converts dU to uracil in a phosphate-dependent reaction. Recombinant DeoA reproduced this activity *in vitro*, and enzymatic depletion of dU attenuated invasive hyphal growth and lesion development. Appressorium-specific expression of *deoA* in *P. oryzae* likewise reduced pathogenicity. Together, these results identify extracellular dU as a factor promoting infection-related development in *P. oryzae* and suggest that dU degradation provides a potential approach for the biological control of rice blast disease.

**IMPORTANCE:** Successful infection by the rice blast fungus *Pyricularia oryzae* depends on tightly regulated developmental changes on the plant surface. This study identifies 2′-deoxyuridine as an extracellular molecule that helps drive these early infection events. The fungus-derived nucleoside promoted appressorium formation and maturation, accumulated during early infection-related development, and could be targeted for disease suppression. A rice field bacterium, *Enterobacter* sp. strain C3, and its enzyme DeoA efficiently degraded 2′-deoxyuridine, and this depletion reduced fungal invasion and disease development. These findings uncover a previously unrecognized extracellular signal associated with infection-related development in the rice blast fungus and point to metabolite degradation by environmental microbes as a promising route for biological control.

## INTRODUCTION

Rice blast, caused by *Pyricularia oryzae* (syn. *Magnaporthe oryzae*), is one of the most destructive diseases of cultivated rice worldwide, causing global yield losses of 10–30% (1). Successful infection by *P. oryzae* depends on a highly coordinated developmental program on the leaf surface. After conidial attachment and germination, the fungus differentiates a specialized infection structure, the appressorium, within a few hours. The appressorium is a hallmark of rice blast pathogenesis and functions as a mechanical device for host invasion (2). As the appressorium matures, glycerol accumulates within the cell, generating substantial turgor pressure that can reach up to 8 MPa. This extreme pressure enables the fungus to breach the rigid rice cuticle by forming a narrow penetration peg that invades host epidermal cells (3). Once inside host tissues, invasive hyphae spread into neighboring cells, initially maintaining a biotrophic interaction before switching to destructive colonization. Over time, expanding lesions develop at the infection site, leading to the production of sporophores and conidia that facilitate rapid disease dissemination (2, 4).

During infection, *P. oryzae* secretes various factors that contribute to host invasion, including effectors and low-molecular-weight metabolites. Although several phytotoxins produced by the rice blast fungus have been described (5), their contributions to early infection-related development remain incompletely understood. In addition to toxins, small extracellular metabolites may influence fungal differentiation and pathogenicity; however, their functional significance during rice blast infection has been only partially characterized.

Previous studies demonstrated that the supernatant of conidial suspensions exhibits pathogenesis-related activity that enhances *P. oryzae* invasion on rice leaves (6–7). The active compound was identified as 2’-deoxyuridine (dU), a pyrimidine 2’-deoxyribonucleoside containing uracil as its nucleobase. dU was shown to function as a self-secreted factor that promotes infection during rice invasion. In leaf-sheath assays, conidia suspended in dU caused enhanced disease development in a dose-dependent manner (7). However, the mechanism by which extracellular dU enhances pathogenicity remains unclear. In particular, it is not known whether dU acts on specific steps of early infection development, whether it accumulates during early infection-related development under conditions conducive to appressorium formation, or whether its activity is influenced by other microorganisms associated with rice plants and cultivation fields.

In this study, we investigated the role of extracellular dU in regulating infection development in *P. oryzae*. We found that dU accelerates early appressorium initiation and enhances appressorium maturation through glycogen mobilization and turgor generation. We also detected extracellular dU during early infection-related development. Based on these observations, we hypothesized that environmental modulation of dU availability could influence disease progression. To test this possibility, dU-degrading bacterial strains were isolated from rice cultivation fields (8). From the strain exhibiting the highest dU-degrading activity, *Enterobacter* sp. strain C3, we identified the responsible dU-metabolizing enzyme as thymidine phosphorylase (deoA) and examined the impact of dU depletion on the pathogenicity of *P. oryzae*. Together, these findings reveal that dU-mediated infection is subject to microbial turnover, highlighting an ecological layer of pathogen regulation during rice blast disease.

## RESULTS

### 2′-Deoxyuridine (dU) accelerates early appressorium initiation without affecting conidial germination

To determine which stage of early infection development in *P. oryzae* is influenced by dU, conidial germination and appressorium formation were quantified on hydrophobic coverslips under different dU concentrations. Conidial germination rates at 3 h were not significantly affected by dU treatment (Fig. 1A). In contrast, appressorium formation at 4 h increased in a dose-dependent manner, with 5 µM dU showing a significant enhancement compared with the control (Fig. 1B). These results indicate that dU does not promote the initial germination of conidia, but rather accelerates the transition from germ tube emergence to appressorium initiation.

**FIG 1.**
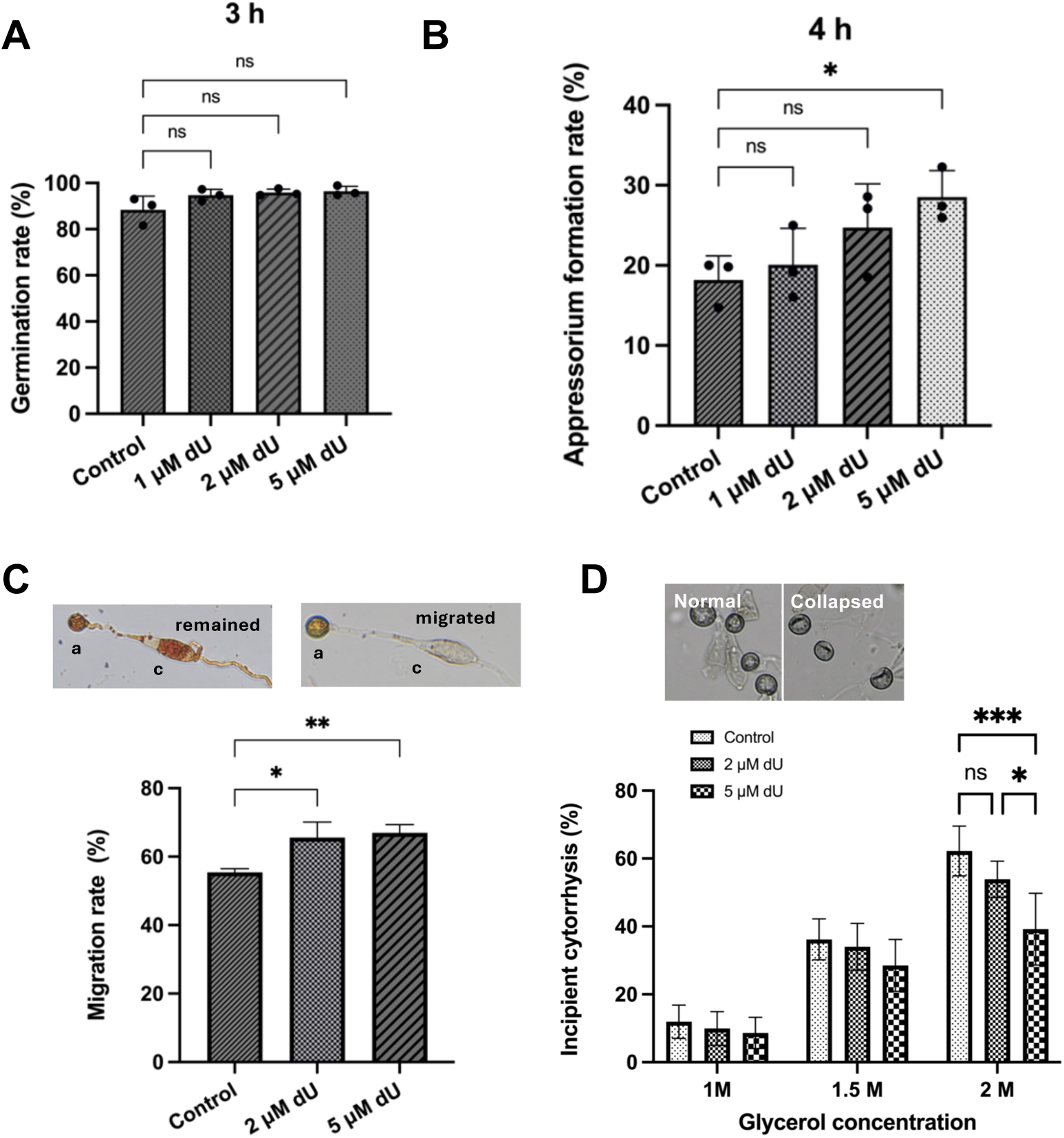
2′-Deoxyuridine (dU) accelerates appressorium initiation and promotes appressorium maturation in *Pyricularia oryzae*. (A) Conidial germination rates at 3 h on hydrophobic coverslips in the presence of the indicated concentrations of dU. No significant differences were detected among treatments. (B) Appressorium formation rates at 4 h on hydrophobic coverslips in the presence of the indicated concentrations of dU. (C) Glycogen mobilization during appressorium maturation at 24 h in the presence of the indicated concentrations of dU. Representative images of conidia retaining glycogen (remained) and conidia showing glycogen migration into the developing appressorium (migrated) are shown above. Quantification represents the percentage of appressoria exhibiting glycogen migration. (D) Intracellular turgor of appressoria at 24 h, assessed by an incipient cytorrhysis assay after exposure to the indicated concentrations of glycerol. Representative images of normal and collapsed appressoria are shown above. Quantification represents the percentage of collapsed appressoria. Data represent means ± SD from three independent biological replicates. Statistical significance in (A–C) was evaluated by one-way ANOVA with Tukey’s post-hoc test. In (D), significance was evaluated by two-way ANOVA followed by Tukey’s multiple comparisons test. ns, not significant; \**P* < 0.05; \*\**P* < 0.01; \*\*\**P* < 0.001.

Consistent with this interpretation, dU had little effect on vegetative growth on solid medium and only modest effects in liquid culture at low concentrations (Fig. S1), suggesting that its primary role is not nutritional but is associated with infection-stage development.

### dU enhances appressorium maturation by promoting glycogen mobilization and turgor generation

Given that functional appressorium maturation depends on the mobilization of stored carbon reserves from the conidium to the developing appressorium, the effect of dU on glycogen dynamics was examined. Glycogen mobilization was quantified 24 h after incubation under different dU concentrations by scoring the proportion of appressoria showing glycogen migration. In the control, 55% of appressoria exhibited glycogen migration, whereas this value increased to 67% in the presence of 5 µM dU (Fig. 1C). These results indicate that dU promotes glycogen mobilization during appressorium maturation.

To determine whether this effect was associated with enhanced turgor generation, a key feature of mature and functional appressoria, intracellular turgor was assessed 24 h after incubation on hydrophobic coverslips using an incipient cytorrhysis assay, in which appressorial collapse was induced by exposure to external glycerol (3). In control samples, 62% of appressoria collapsed upon exposure to 2 M glycerol, whereas this proportion was reduced to about 40% following treatment with 5 µM dU, indicating elevated intracellular turgor in dU-treated appressoria (Fig. 1D). In addition, appressoria exposed to 5 µM dU exhibited significantly larger diameters than untreated controls, whereas no significant difference was detected at 2 µM dU (Fig. S2). Together, these results indicate that dU enhances appressorium maturation by promoting glycogen mobilization, turgor generation, and associated morphological development.

### Extracellular dU accumulates during early infection-related development in *P. oryzae* conidial suspensions

To determine whether extracellular dU accumulates during early infection-related development, conidial suspensions of *P. oryzae* at different densities were inoculated onto barley leaves, and extracellular fluids were collected after 4 h and 20 h for HPLC analysis. At 4 h, when appressorium formation had begun, extracellular dU was detected at all tested conidial densities and increased from 1 × 10^5^ to 1 × 10^6^ conidia/mL. However, dU did not increase further at 2 × 10^6^ conidia/mL (Fig. 2). At 20 h, when appressoria were largely mature, extracellular dU concentrations showed a different pattern depending on the initial conidial density and reached their highest level at 1 × 10^6^ conidia/mL (Fig. 2). These results indicate that extracellular dU accumulates during early infection-related development and that its abundance depends on both incubation time and conidial density.

**FIG 2.**
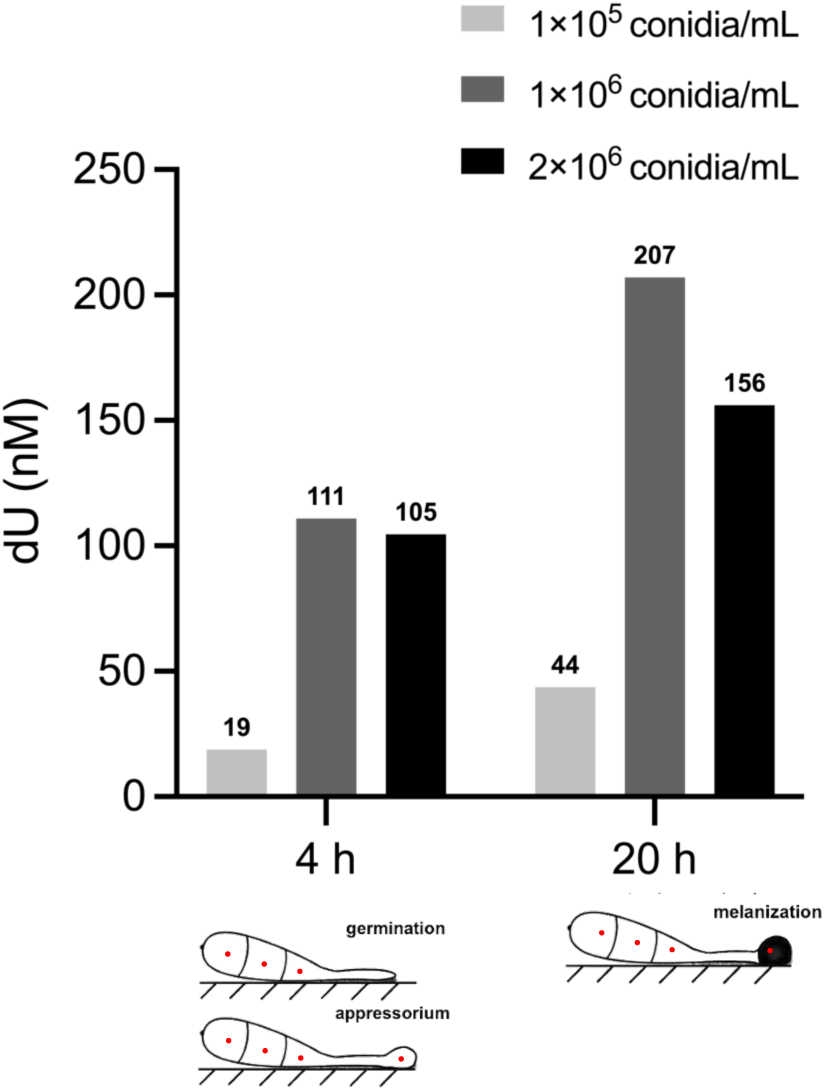
Effect of *Pyricularia oryzae* conidial density on extracellular dU accumulation during early infection-related development. Conidial suspensions of *P. oryzae* at different densities were inoculated onto barley leaves and incubated for 4 h or 20 h. Extracellular fluids were then collected and analyzed for dU by HPLC. The 4-h time point corresponds to the onset of appressorium initiation, whereas the 20-h time point corresponds to largely mature appressoria. The schematic summarizes the relationship between conidial density and extracellular dU accumulation during these stages.

#### Isolation of dU-metabolizing bacteria and identification of the metabolic enzyme

Given that dU enhances the pathogenicity of rice blast disease, microorganisms capable of degrading dU were isolated from rice field environments. More than 120 bacterial strains were obtained from soil, leaves, leaf sheaths, and rhizosphere samples of rice (8). Their dU-degrading activities were compared, and strain C3 was selected because it showed the most rapid dU degradation. Based on 16S rRNA sequence analysis and whole-genome sequencing, strain C3 was assigned to the genus *Enterobacter* (8).

To determine the metabolite produced from dU by strain C3, reaction products were first analyzed by TLC together with authentic standards. The product spot co-migrated with uracil, indicating that uracil was the major dU-derived metabolite (Fig. 3A). The TLC-separated product was then recovered from the plate and analyzed by GC-MS after trimethylsilyl (TMS) derivatization. The resulting spectrum was consistent with uracil, further supporting the identification of uracil as the metabolite produced from dU by strain C3 (Fig. 3B). In addition, the crude enzyme extract showed high activity in potassium phosphate (KPi) buffer but much lower activity in Tris-HCl buffer (Fig. S3), indicating phosphate dependence of the reaction. Because the reaction was phosphate-dependent and yielded uracil as the major product, the responsible enzyme was inferred to be a nucleoside phosphorylase.

**FIG 3.**
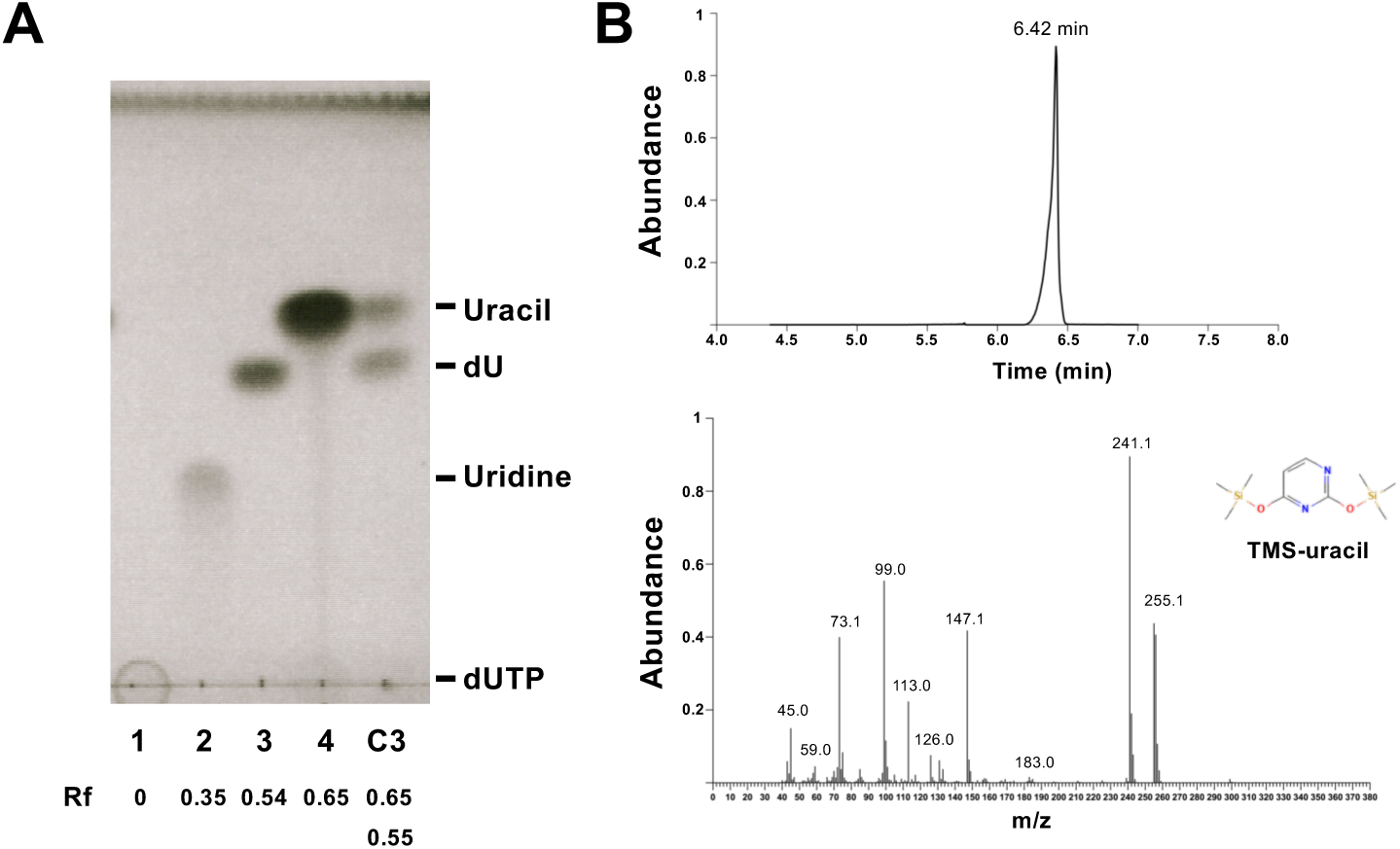
Identification of the dU-derived metabolite produced by *Enterobacter* sp. strain C3. (A) TLC analysis of metabolites produced from dU by strain C3 after 1 h of incubation, shown alongside authentic standards. After incubation of dU with strain C3, the dU spot was reduced, and a new spot appeared that co-migrated with authentic uracil. An additional spot was also detected. Rf values are indicated in the figure. Lane 1, dUTP; lane 2, uridine; lane 3, dU; lane 4, uracil. (B) GC-MS analysis of the TLC-isolated product after trimethylsilyl (TMS) derivatization. The spectrum was consistent with TMS-uracil [2,4-bis(trimethylsilyloxy)pyrimidine], supporting the identification of uracil as a metabolite produced from dU by strain C3.

To identify the enzyme responsible for dU metabolism, bacterial cell extracts and culture supernatants of strain C3 were first assayed for activity. The highest activity was detected in the crude intracellular fraction (Fig. S3). The active protein was then enriched from bacterial lysates by sequential chromatography, including DEAE-Sepharose anion-exchange chromatography, Butyl-650M hydrophobic interaction chromatography (HIC), and size-exclusion chromatography (SEC). In the final SEC step, dU-metabolizing activity was associated with a distinct peak corresponding to an estimated native molecular mass of approximately 50 kDa (Fig. 4A). SDS-PAGE of the active fractions revealed a protein band near 50 kDa whose abundance pattern broadly matched the activity profile across fractions (Fig. 4B). To identify the enzyme, selected fractions from the final SEC step were further compared by nanoLC–MS/MS analysis using a Q Exactive Orbitrap mass spectrometer. Among the fractions examined, fractions 8, 11, and 14 were chosen as representative fractions with low, high, and intermediate dU-metabolizing activity, respectively, to compare their protein profiles and search for candidates involved in dU metabolism. Initial analysis yielded multiple candidate proteins with similar predicted molecular masses, making it difficult to assign the activity to a single protein.

**FIG 4.**
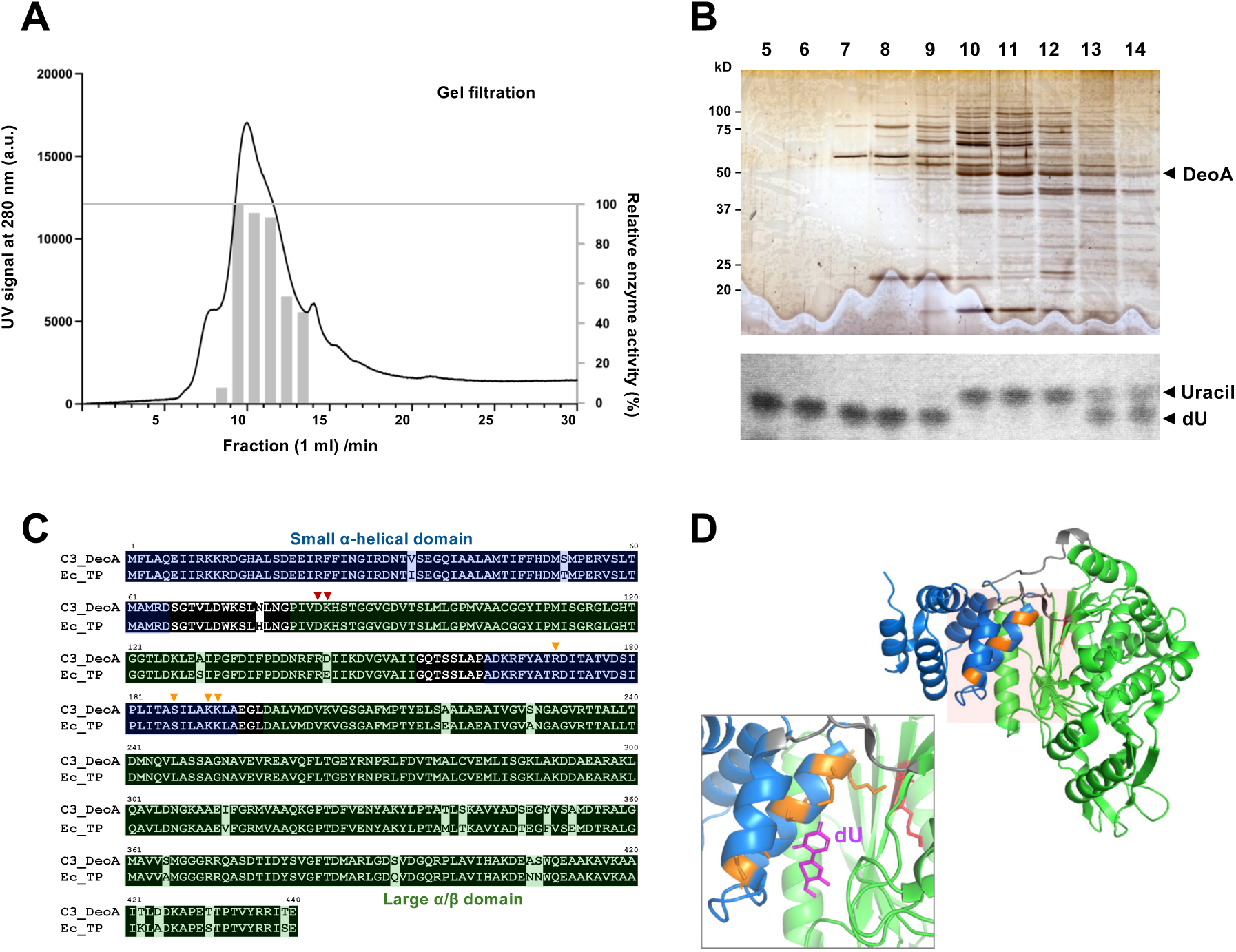
Identification of DeoA as the dU-metabolizing enzyme from *Enterobacter* sp. strain C3. (A) Size-exclusion chromatography (SEC) elution profile of the strain C3 extract. Grey bars indicate fractions positive for dU-metabolizing activity. (B) SDS-PAGE analysis of the indicated SEC fractions (top) and the corresponding TLC-based assay for dU-metabolizing activity (bottom). Fraction numbers are shown above the lanes. A protein band of approximately 50 kDa (arrowhead, DeoA) was enriched in fractions showing dU-metabolizing activity. Positions of uracil and dU in the TLC assay are indicated on the right. (C) Sequence alignment of the C3 DeoA candidate with *Escherichia coli* thymidine phosphorylase (Ec TP). The small α-helical domain and large α/β domain are highlighted in blue and green, respectively. Conserved catalytic residues are indicated by arrowheads. (D) AlphaFold-predicted structure of the C3 DeoA candidate. The small α-helical domain and large α/β domain are shown in blue and green, respectively. Conserved catalytic residues are highlighted in red and orange. Insets show a close-up of the putative active-site region and the placement of dU (magenta) relative to the homologous ligand-bound *E. coli* thymidine phosphorylase structure (14).

We therefore re-evaluated the dataset using three criteria: consistency with the estimated molecular mass from SEC and SDS-PAGE, correlation between protein abundance across fractions and the activity profile, and predicted enzymatic function consistent with the phosphate-dependent conversion of dU to uracil. Candidate proteins were prioritized primarily based on peptide-spectrum matches (PSMs) and relative abundance in each fraction. This integrative comparison identified DeoA as the most plausible candidate for the dU-metabolizing enzyme.

Because the draft genome sequence of strain C3 had been obtained (8), the sequences of candidate proteins could be retrieved and further examined by sequence alignment and structure prediction. This analysis identified a single strong candidate, thymidine phosphorylase (DeoA), with a predicted molecular mass of approximately 49 kDa, consistent with the size estimated by gel filtration and SDS-PAGE (Fig. 4A, B). Sequence alignment and structure prediction further supported the assignment of this protein as a DeoA-like nucleoside phosphorylase (Fig. 4C, D).

A phylogenetic analysis of DeoA-like proteins further indicated that closely related proteins are broadly distributed across members of the Enterobacterales (Fig. S4). The C3 DeoA sequence clustered with homologs from several enterobacterial taxa, including *Citrobacter*, *Leclercia*, and *Cedecea*, suggesting that DeoA-like dU-metabolizing enzymes are not unique to strain C3 but may be more widely conserved among enterobacterial bacteria.

### Enzymatic depletion of dU attenuates the pathogenicity of *P. oryzae*

To determine whether DeoA indeed possesses dU-metabolizing activity, *deoA* was cloned into the pET-30a vector, expressed in *E. coli* BL21, and purified as a recombinant 6×His-DeoA protein. Purified DeoA showed clear dU-metabolizing activity in TLC-based assays (Fig. 5A, B). As observed for the crude extract of strain C3 (Fig. S3), the activity was markedly lower in Tris-HCl (pH 7) than in potassium phosphate (KPi) buffer (pH 7), indicating phosphate dependence of the recombinant enzyme (Fig. 5B).

**FIG 5.**
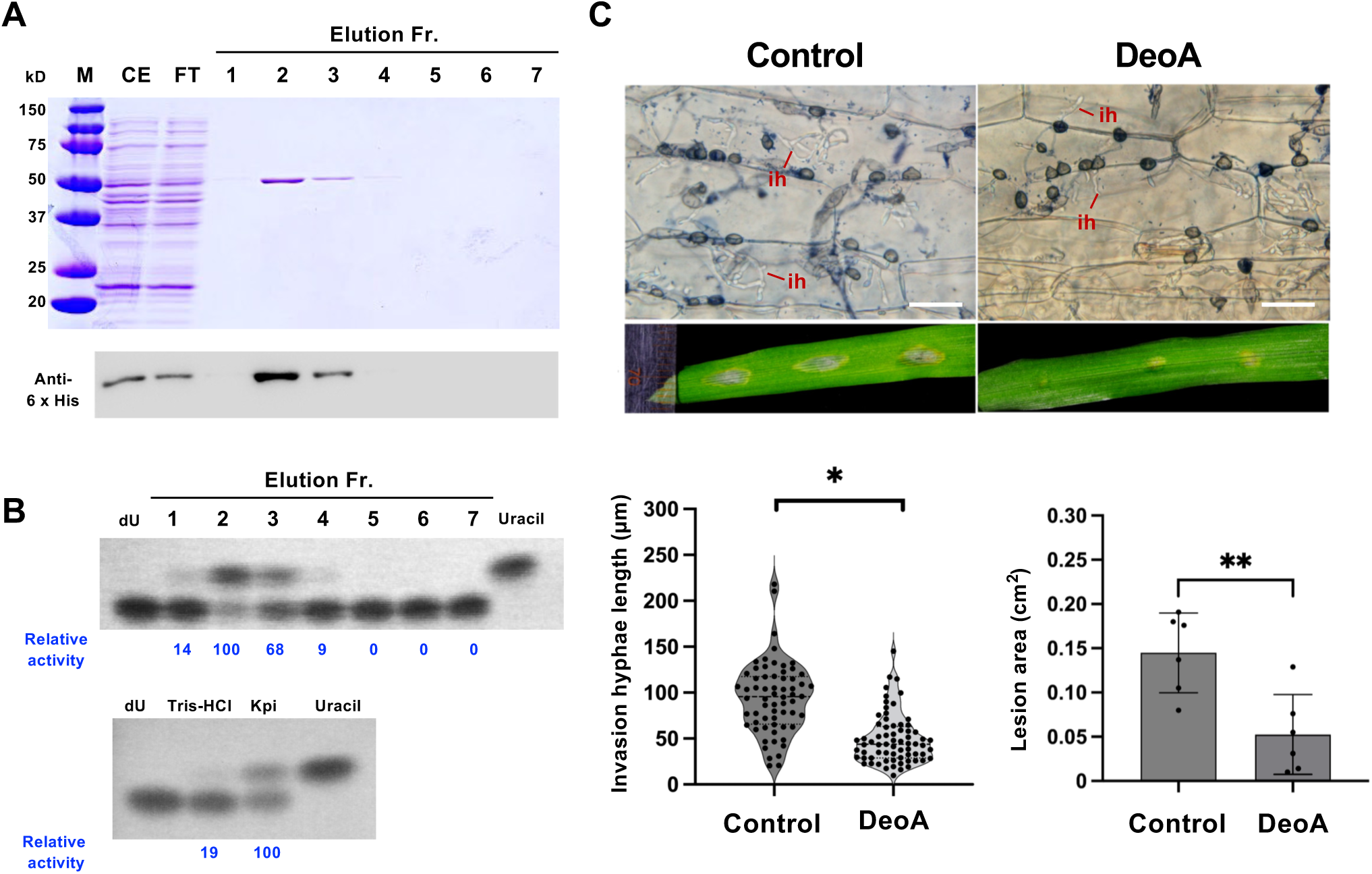
Heterologous expression of DeoA from *Enterobacter* sp. strain C3 in *E. coli* and examination of its effects on *P. oryzae* infection. (A) Purification of recombinant 6×His-DeoA heterologously expressed in *E. coli* by Ni-affinity chromatography, shown by SDS-PAGE (upper) and anti-6×His immunoblotting (lower). M, molecular mass markers; CE, cell extract; FT, flow-through; 1–7, elution fractions. (B) TLC-based assay of dU-metabolizing activity using the purified fractions shown in (A) (upper) and using purified DeoA (Fr. 2) under different buffer conditions (lower). Relative activity values, normalized to the most active sample, are shown below the TLC panels. (C) The effects of recombinant DeoA on *P. oryzae* infection were examined using barley sheath and detached-leaf assays. Representative microscopic images of invasive hyphae (ih) in barley sheath cells are shown in the upper panels, and representative lesion symptoms on barley leaves are shown in the middle panels. The lower panels show quantification of invasive hyphal length and lesion area in control and DeoA-treated samples. Each dot represents an individual measurement. Data represent means ± SD from independent biological replicates. Statistical significance was evaluated using a two-tailed Student’s *t*-test. * *P* < 0.05; ** *P* < 0.01.

Next, fresh conidia were mixed with purified DeoA and inoculated onto barley leaves. Compared with the control, DeoA-treated samples showed reduced disease symptoms and significantly restricted invasive hyphal growth (Fig. 5C). These results indicate that enzymatic depletion of extracellular dU attenuates rice blast infection.

### Appressorium-specific expression of deoA compromises the pathogenicity of *P. oryzae*

To investigate the *in vivo* functional significance of *deoA*, we attempted to express it in *P. oryzae*. However, repeated attempts driven by the constitutive TEF promoter failed to yield viable transformants, suggesting that continuous expression of *deoA* may be detrimental during early developmental stages. To circumvent this problem, we employed the appressorium-specific GAS2 (MGG_04202) promoter, which is active during appressorium formation but inactive during vegetative growth (9). Using this stage-specific expression system, DeoA-GFP-positive transformants were obtained. Fluorescence microscopy confirmed that DeoA-GFP accumulated specifically in differentiating appressoria on plastic coverslips (Fig. 6A). Although the DeoA-GFP transformants formed appressoria normally, they exhibited reduced virulence on barley leaves, and produced smaller lesions than the control transformant expressing GFP (Fig. 6B). These results indicate that appressorium-specific expression of *deoA* compromises the pathogenicity of *P. oryzae*.

**FIG 6.**
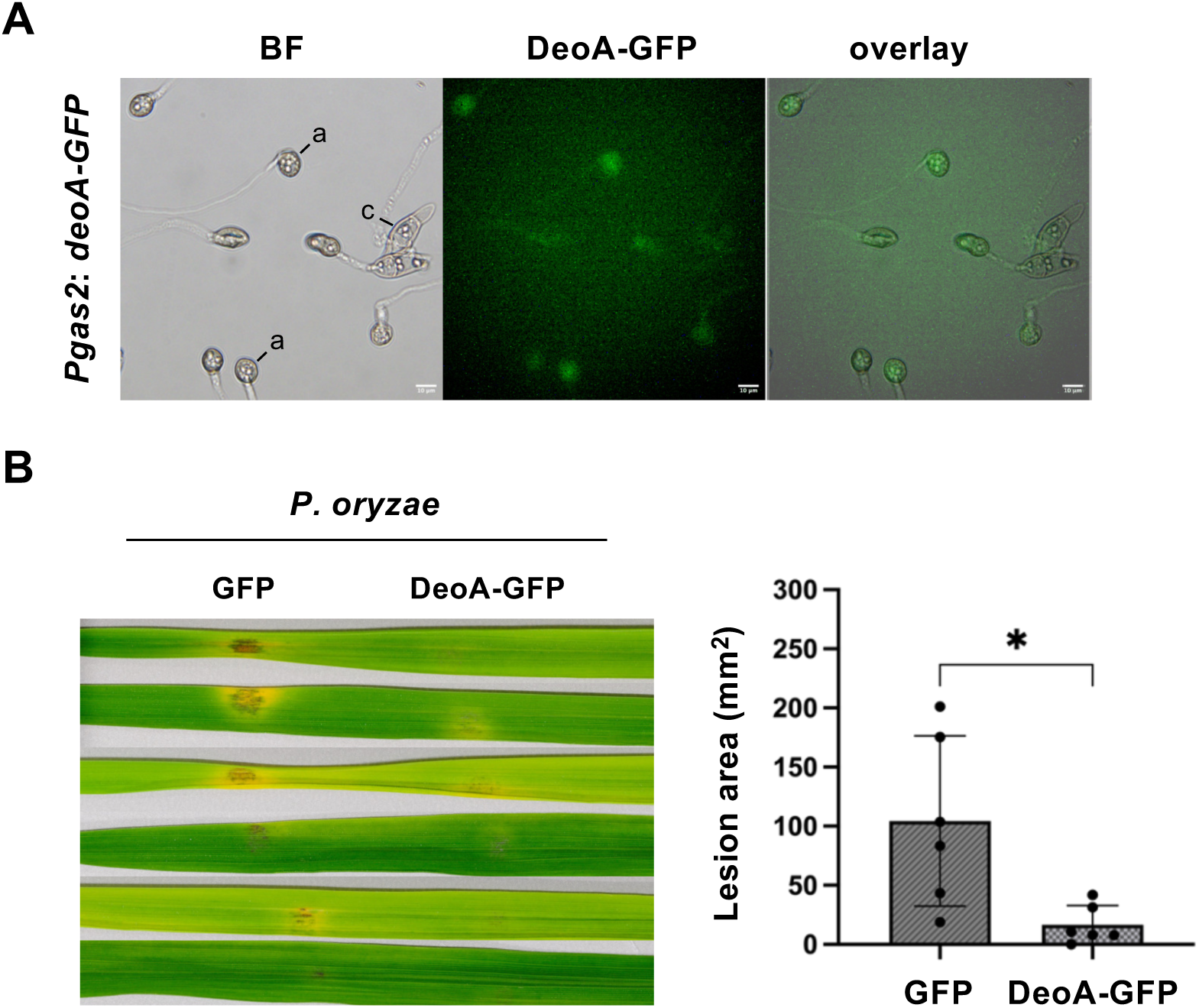
Appressorium-specific expression of deoA compromises the pathogenicity of *P. oryzae*. (A) Bright-field (BF), fluorescence, and merged images of a *P. oryzae* transformant expressing DeoA-GFP under the control of the appressorium-specific *GAS2* promoter. GFP fluorescence was detected specifically in differentiating appressoria on plastic coverslips. a, appressorium; c, conidium. Bars = 10 µm. (B) Disease symptoms on barley leaves inoculated with *P. oryzae* transformants expressing GFP under the TEF promoter or DeoA-GFP under the GAS2 promoter, with lesion area quantified at right. Individual data points are shown; bars represent means ± SD. Statistical significance was evaluated using a two-tailed Student’s *t*-test. * *P* < 0.05.

## DISCUSSION

In this study, extracellular dU was found to promote infection-related development in *P. oryzae* and to contribute to fungal pathogenicity. Although dU had limited and condition-dependent effects on vegetative growth, it more clearly enhanced early appressorium formation, glycogen mobilization, and turgor generation, all of which are important for successful host penetration. Extracellular dU was detected during early infection-related development, supporting the idea that dU is present under conditions relevant to appressorium formation and early surface-associated development. Furthermore, enzymatic depletion of dU by the bacterial nucleoside phosphorylase DeoA attenuated fungal infection, indicating that extracellular dU is not merely correlated with infection but functions as a metabolite that promotes disease development.

Previous studies showed that exogenous dU enhances lesion development by *P. oryzae* on detached rice leaves. However, dU neither caused visible damage to rice plants nor suppressed host defense responses, including the expression of pathogenesis-related (PR) genes, the production of H_2_O_2_ and phytoalexins (6–7). These observations suggested that the infection-promoting effect of dU is mediated primarily through the fungal side rather than through direct suppression of host immunity. Consistent with this interpretation, the present results indicate that dU acts not on conidial germination itself, but more specifically on appressorium initiation and maturation. Both appressorium formation and intracellular turgor increased in a dose-dependent manner; that is, higher dU concentrations (5 µM) led to enhanced early appressorium formation and raised turgor pressure. Glycogen mobilization during appressorium development, which is crucial for turgor generation (10–11), was also promoted by dU, suggesting that dU facilitates appressorium maturation in addition to appressorium initiation. This interpretation is further supported by the reduced frequency of collapse in the incipient cytorrhysis assay (12). Together, these findings suggest that dU primarily promotes the developmental transition from germ tube emergence to a functional appressorium, thereby facilitating successful host penetration.

Because dU is a common nucleotide metabolite, its possible involvement in the early stages of infection was examined. Extracellular dU concentration increased with conidial density, although this effect did not increase further at the highest conidial density tested (Fig. 2). Extracellular dU was detected in the nanomolar range during early infection development and increased further at the later time point, indicating that dU accumulates during early infection-related development in a manner dependent on both conidial density and incubation time (Fig. 2). This trend is broadly consistent with previous studies showing that dU concentration can influence infection-related outcomes on rice leaves (7). Although the dU concentrations detected in bulk extracellular fluids were lower than the micromolar concentrations used in some *in vitro* assays, dU may become more concentrated in localized microenvironments associated with surface moisture, such as small water droplets formed under humid conditions (13). Such local enrichment may help explain how extracellular dU influences fungal development during early infection.

The identification of *Enterobacter* sp. strain C3 and its dU-metabolizing enzyme DeoA provides complementary support for the functional importance of extracellular dU during infection. Strain C3 converted dU to uracil, and both the crude extract and purified recombinant DeoA showed markedly higher activity in potassium phosphate buffer than in Tris-HCl buffer, consistent with a phosphate-dependent nucleoside phosphorylase reaction (14). This biochemical behavior, together with the activity-guided purification, predicted molecular mass, and structural similarity to thymidine phosphorylase (15), supports the assignment of DeoA as the enzyme responsible for dU metabolism. Importantly, the infection-promoting effect of dU could be counteracted by enzymatic depletion: purified recombinant DeoA attenuated invasive hyphal growth and lesion development, and appressorium-specific expression of deoA in *P. oryzae* likewise reduced pathogenicity. These findings strengthen the view that extracellular dU is functionally required for full infection rather than being a passive by-product of fungal growth.

This metabolite might act as a localized infection-promoting signal rather than a classical QS molecule (16–17). More specifically, dU may function as a developmental cue that fine-tunes fungal behavior according to its local concentration and microenvironmental context on the leaf surface during early invasion. In this sense, extracellular dU appears to be better understood as a site-restricted signal associated with infection-stage development than as a canonical quorum-sensing molecule. From an applied perspective, the identification of dU-degrading bacteria and enzymes suggests a potential new strategy for disease control. Microorganisms such as strain C3, or enzymes such as DeoA, may help reduce disease development by lowering extracellular dU during early infection-related development. Further studies will be needed to determine how broadly this mechanism operates under natural conditions and whether dU-scavenging activity in the phyllosphere microbiota contributes to disease suppression (18–19).

## MATERIALS AND METHODS

### Biological materials, strains, media, and chemicals

Rice (*Oryza sativa* L. cv. Koshihikari) and barley (*Hordeum vulgare* L. cv. Syunrai) were used in this study. Plants were grown in soil-filled pots at 26°C under a 16-h-light/8-h-dark cycle. The soil (Kumiai Nippi Engei Baido No. 1; Nippi, Inc., Tokyo, Japan) was autoclaved at 121°C for 30 min before use. Two-week-old barley and four-week-old rice plants were used for detached-leaf assays. The fungal isolates used in this study were *Pyricularia oryzae* Ken53-33 (MAFF 101525, race 007.0) and Ina86-137 (MAFF 101511, race 007.0). 2′-Deoxyuridine (dU) used for incubation experiments and HPLC standardization was purchased from Wako Pure Chemical Industries (Osaka, Japan).

### Conidial germination, appressorium formation, glycogen staining, and appressorial turgor and size measurements

To investigate the infection-promoting mechanism of dU in *P. oryzae*, the effects of dU treatment on conidial development on a hydrophobic surface were examined. Plastic coverslips (Thermo Fisher Scientific) were sterilized by UV irradiation for at least 30 min. Conidial suspensions were prepared at a final concentration of 1 × 10^5^ conidia/mL. To induce appressorial development, six to nine 10-µL droplets were placed on each coverslip and incubated in a humid chamber at 28°C. Conidial germination was quantified after 3 h, and appressorium formation was quantified after 4 h. For glycogen staining and turgor measurements, samples were incubated for 24 h. More than 100 appressoria were counted on each coverslip. Glycogen was visualized by mounting the coverslips in a staining solution containing 60 mg KI and 10 mg I_2_ per mL distilled water. Appressorial turgor was assessed by an incipient cytorrhysis assay. After 24 h of incubation, water was carefully removed from the coverslips, and an equal volume of 0.5, 1.0, or 2.0 M glycerol was added. The number of collapsed appressoria was recorded under a microscope after 10 min. For appressorial size measurements, images were captured at 400× magnification after 24 h. Images were converted to 8-bit format, and the threshold was adjusted until the dark appressorial structure could be clearly identified. Appressorial diameter was then measured using the Hough Circle Transform plugin in ImageJ (version 2.14.0/1.54f).

### Vegetative growth assay

Complete medium (CM, 20) was used for *P. oryzae* growth. For solid-medium growth assays, *P. oryzae* was cultured on CM agar at 28°C for 11 days in the presence or absence of dU, and colony area was measured using ImageJ. For liquid culture assays, 100 µL of conidial suspension (1 × 10^6^ conidia/mL) was inoculated into liquid CM containing 0, 1, or 5 µM dU and incubated at 28°C with shaking at 120 rpm for 6 days. Mycelia were collected by filtration, washed twice with distilled water, gently blotted dry with absorbent paper, and transferred to weighing paper. The samples were then dried to constant mass in a forced-air oven at 80°C. Biomass measurements were performed in three independent biological replicates.

### Quantification of extracellular dU during early infection-related development

To quantify extracellular dU accumulation during early infection-related development, conidial suspensions of *P. oryzae* at the indicated densities were inoculated onto barley leaves and incubated in a humidified chamber at 28°C. Extracellular fluids were collected at 4 h and 20 h after incubation. To remove conidia and other debris, the collected liquid was centrifuged at 20,000 × g for 10 min, and the resulting supernatant was lyophilized and resuspended in distilled water. Prior to HPLC analysis, a 200-µL aliquot of each resuspended sample was filtered through a 0.45-µm membrane filter using a 1-mL syringe and transferred into a vial insert. Extracellular dU was quantified using a Shimadzu HPLC system (Shimadzu, Kyoto, Japan) equipped with a Develosil ODS column (Nomura Chemical, Seto, Japan). Chromatographic separation was performed with an isocratic mobile phase consisting of 95% Milli-Q water and 5% acetonitrile at a total flow rate of 1.0 mL/min. The column temperature was maintained at 40°C, and the injection volume was 10 µL. Analytes were monitored using a photodiode array detector, and the total run time was 15 min per sample. dU concentrations were determined based on comparison with an authentic dU standard.

### Isolation and screening of dU-degrading bacteria

Isolation and screening of dU-degrading bacteria were performed essentially as described previously (8). Briefly, environmental samples collected from rice field environments, including bulk soil, rhizosphere soil, leaves, roots, and leaf sheaths, were inoculated into dU minimal medium (dUMM; composition in Table S1), in which 2′-deoxyuridine was supplied as the sole carbon source. Cultures showing dU-degrading activity were serially diluted and plated onto Reasoner’s 2A (R2A, 21) agar to isolate single colonies. Purified isolates were cryopreserved in glycerol at − 80°C and compared for dU-degrading activity after incubation in dUMM at 28°C for 24 h. The isolate showing the highest activity was selected and designated strain C3. To further purify the target bacterium, suspensions from the second enrichment culture were streaked onto MacConkey agar, and pink-to-red colonies were selected. Taxonomic assignment of strain C3 was performed by 16S rRNA gene sequencing using the universal bacterial primers 10f_16S and 1541r_16S, followed by BLASTN analysis against the NCBI nucleotide database.

### Purification of the dU-metabolizing enzyme from *Enterobacter* sp. strain C3

*Enterobacter* sp. strain C3 was cultured in 1 L of Luria-Bertani (LB) broth at 28°C with reciprocal shaking at 150 rpm for 48 h. Cells were harvested by centrifugation (7,000 × g, 15 min, 10°C), washed, and resuspended in 20 mM potassium phosphate buffer (pH 7.0). The cell suspension was lysed by sonication on ice, and cell disruption efficiency was verified microscopically. The lysate was clarified by centrifugation (20,000 × g, 15 min, 4°C) and filtered through a 0.45-µm membrane.

To reduce non-specific binding, the crude extract was supplemented with 50 mM NaCl and applied to a DEAE-Sepharose CL-6B anion-exchange column equilibrated with 20 mM potassium phosphate buffer (pH 7.5) containing 10% glycerol and 0.1 mM DTT. Proteins were eluted with a linear NaCl gradient (50–500 mM), and fractions showing dU-metabolizing activity were pooled and concentrated by ultrafiltration (10,000 MWCO). The concentrated sample was adjusted to 20% ammonium sulfate saturation and loaded onto a TOYOPEARL Butyl-650M hydrophobic interaction chromatography column. Proteins were eluted using a descending linear gradient of ammonium sulfate (20% to 5%) in the same buffer system. Active fractions were collected, concentrated, and further purified by size-exclusion chromatography using a TSK-GEL G3000SW column connected to an HPLC system (Shimadzu SPD-M20A detector). The column was operated at 10°C with 20 mM potassium phosphate containing 0.1 M NaCl (pH 7.0) at a flow rate of 0.8 mL/min, with an injection volume of 100 µL. The native molecular weight of the enzyme was estimated by calibration with standard proteins: bovine serum albumin (67 kDa), ovalbumin (43 kDa), chymotrypsinogen A (25 kDa) and ribonuclease A (13.7 kDa) (Fig. S5).

### TLC assay and GC-MS identification of the dU-derived metabolite

To identify the metabolite produced from dU by *Enterobacter* sp. strain C3, reaction products were first analyzed by thin-layer chromatography (TLC). Aliquots (2 µL) of the reaction mixture were spotted onto aluminum-backed silica gel 60 F254 plates (Merck). The plates were developed at room temperature in chloroform:methanol (4:1, v/v) for approximately 10 min, air-dried, and visualized under UV light (254 nm). Authentic standards were analyzed in parallel. For identification of the dU-derived metabolite, the corresponding TLC spot was recovered from the plate and subjected to GC-MS analysis after trimethylsilyl (TMS) derivatization. Gas chromatography was performed on an Agilent Technologies 7890A GC System using a DuraBond Ultra Inert column (length 30 m; diameter 0.25 mm; film 0.25 µm, Agilent Technologies, USA) as previously described (22). The resulting mass spectrum was compared with that of the authentic compound to determine the identity of the metabolite.

### SDS-PAGE and immunoblotting

Proteins were boiled in 5×loading buffer (with 5% β-mercaptoethanol) and separated on 12% SDS-PAGE gels (140 V, 90 min; Biocraft, Japan) before CBB or silver staining (Bio-Rad, USA). For western blotting, proteins were semi-dry transferred (ATTO, Japan) to 0.45-µm nitrocellulose membranes. Membranes were blocked (3% milk-TBS, 1 h) and probed with HRP-conjugated anti-6×His antibody (1:1,000; Wako, Japan) for 30 min. Following washes in TBS-T, protein bands were detected using Western BLoT HRP Substrate (Takara, Japan) and a LuminoGraph II imager (ATTO, Japan).

### NanoLC-MS/MS protein identification

Selected fractions (fractions 8, 11, and 14) from the final size-exclusion chromatography step were analyzed by nanoLC-MS/MS using a Q Exactive Orbitrap mass spectrometer (Thermo Fisher Scientific, Waltham, MA, USA). Samples were prepared using an SP3-based digestion method. Raw data were processed with Proteome Discoverer (version 2.4.1.15; Thermo Fisher Scientific) using Sequest HT and Percolator and searched against a UniProt database containing *Enterobacter asburiae* protein sequences with a false discovery rate (FDR) of ≤1%. Proteins detected in the selected fractions were compared primarily on the basis of peptide-spectrum matches (PSMs) and abundance values.

### AlphaFold-based structural analysis of the DeoA candidate

The predicted structure of the C3 DeoA candidate (UniProt accession A0AAP5VD06) was retrieved from the AlphaFold Protein Structure Database (23) and visualized in PyMOL (Schrödinger, LLC, ver. 3.1.0). Domain boundaries were assigned based on homology to *Escherichia coli* thymidine phosphorylase, and conserved catalytic residues were identified by sequence comparison (15). The ligand-bound structure of *E. coli* thymidine phosphorylase (PDB ID: 4EAD) was superposed onto the AlphaFold model, and the bound ligand (0NP) was used as a positional reference for the putative substrate-binding site. A dU molecule was manually placed in the active-site cleft for schematic comparison.

### Phylogenetic analysis of DeoA-like proteins

DeoA-like proteins were retrieved by BLASTP search against the NCBI Clustered NR database (accessed on March 30, 2026) using the amino acid sequence of *Enterobacter* sp. strain C3 DeoA as the query. The sequences included in the phylogenetic analysis are listed in Table S2. Amino acid sequences were aligned using MAFFT (v7.526, option: --auto; 24). Both untrimmed and trimmed alignments were analyzed; trimming was performed using trimAl (v1.5.rev1, option: -automated1; 25). Maximum-likelihood phylogenetic trees were inferred using IQ-TREE v3.0.1 with ModelFinder (26–27). For the trimmed alignment, the best-fit substitution model selected according to BIC was Q.PFAM+I+R6. Branch support was evaluated with 1,000 ultrafast bootstrap replicates and 1,000 SH-aLRT replicates (28–29). The resulting trees were visualized in FigTree v1.4.4, with the *Enterobacter* sp. strain C3 DeoA sequence highlighted in the final tree.

### Vector construction

For in vitro protein expression, the full-length coding sequence of *deoA* was amplified from C3 genomic DNA. The resulting fragment was cloned into the pET-30a(+) vector (Novagen, Madison, WI, USA), which had been linearized with *Not*I and *Eco*RV, using the In-Fusion HD Cloning Kit (Takara Bio, Japan) to generate pET-30a-deoA. For stage-specific expression in *P. oryzae*, the amplified *deoA* sequence was fused with eGFP and inserted into the pPN94 (30) under the control of the appressorium-specific *GAS2* (MGG_04202) promoter using the In-Fusion HD Cloning Kit (Takara Bio, Japan). Primer sequences used for vector construction are listed in Table S3. All constructs were verified by Sanger sequencing prior to bacterial transformation or fungal protoplast transformation.

### Heterologous expression and purification of recombinant DeoA

The pET-30a-deoA construct was transformed into *E. coli* BL21(DE3) competent cells by electroporation (MicroPulser, Bio-Rad, USA). Transformants were cultured in SOC medium supplemented with 50 µg/mL kanamycin at 37°C until the OD_600_ reached approximately 0.6. Protein expression was induced with 0.5 mM isopropyl-β-D-thiogalactopyranoside (IPTG), followed by incubation at 37°C for 16 h. Cells were harvested by centrifugation, resuspended in lysis buffer (50 mM KPi buffer, pH 7.0, 200 mM NaCl, 20 mM imidazole), and disrupted by sonication on ice. After centrifugation (12,000 × *g*, 15 min, 4°C), the supernatant was passed through a 0.45-µm filter and loaded onto a manually packed 1-mL column with Ni-agarose resin (141-07983; FUJIFILM Wako, Japan). The column was washed with 20 column volumes (CV) of KPi buffer (50 mM imidazole, 200 mM NaCl), and the recombinant protein was eluted stepwise with 1 CV aliquots of KPi buffer containing 100-, 200-, and 500-mM imidazole, respectively. Target fractions, confirmed by SDS-PAGE and immunoblotting, were pooled and buffer-exchanged into 20 mM KPi (pH 7.0) via five cycles of centrifugal ultrafiltration (10-kDa MWCO; Violamo, AS ONE, Japan) to remove residual salt. Protein concentration was determined using the Bio-Rad protein assay (Bio-Rad, Hercules, USA) according to the manufacturer’s instructions.

### Protoplast preparation and transformation of *P. oryzae*

Protoplast preparation and PEG-mediated transformation of *P. oryzae* were performed essentially as previously described (31). with minor modifications. Vegetative mycelia cultured in liquid CM for 3 days were harvested and digested in osmotic buffer (1.2 M MgSO_4_, 10 mM sodium phosphate, pH 5.8) containing 10 mg/mL cellulase (Yakult Pharmaceutical Industry Co., Ltd, Japan) and 5 mg/mL kitalase (FUJIFILM Wako) at 28°C for 3–4 h. The filtrate was slowly overlaid with an equal volume of ice-cold ST buffer and centrifuged using a swing rotor. Undigested cell debris was pelleted at the bottom, while the protoplasts were recovered from the interface. Protoplasts were collected from the interface, washed twice with STC buffer, and resuspended to 1 × 10^8^ cells/mL with STC.

For transformation, 100 µL of protoplast suspension was mixed with 5–10 µg of plasmid DNA and incubated at RT for 20 min. Following the addition of 1 mL PTC, the mixture was incubated at RT for another 20 min. The suspension was then thoroughly mixed with YBSA regeneration medium (equilibrated at 50°C; 1 M sucrose, 0.1% casein hydrolysate, 0.1% yeast extract, 1.8% agar) and plated. After 36 h of incubation at 28°C, the plates were overlaid with CM top agar containing 250 µg/mL hygromycin B. Putative transformants were recovered after 5–7 days, purified by single-spore isolation, and confirmed by PCR and fluorescence microscopy.

### Infection assays and microscopic analyses

The effects of recombinant DeoA on *P. oryzae* infection were examined using barley sheath and detached-leaf assays. Barley detached-leaf assays were performed by inoculating 20 µL droplets of conidial suspensions (1 × 10^5^ conidia/mL) containing 83.3 µg/mL DeoA or buffer control. Leaves were secured in a humidity-controlled chamber and incubated in the dark at 28°C for 6 days, after which lesion development was quantified via ImageJ.

For microscopic analyses, barley sheath tissues inoculated with conidial suspensions were incubated under humid conditions at 28°C for 24 h. The tissues were decolorized in methanol (80 rpm, RT), and then stained with trypan blue to visualize invasive hyphal growth as previously described (32). Stained tissues were examined microscopically, representative images were recorded, and invasive hyphal length was quantified using ImageJ. GFP fluorescence in *P. oryzae* transformants expressing DeoA-GFP was observed using a BX51 fluorescence microscope (Olympus, Tokyo, Japan). Bright-field and fluorescence images were captured under identical settings and merged for figure preparation.

## ACKNOWLEDGMENTS

This work was supported by the Japan Society for the Promotion of Science (JSPS) through Grant-in-Aid for Scientific Research (C) (21K05595) to IS, and through Challenging Research (Exploratory) (22K19176) and Grant-in-Aid for Scientific Research (B) (23K26905) to DT. H.H. was supported by JST SPRING (JPMJSP2125) through the THERS Make New Standards Program for the Next Generation Researchers. ChatGPT (OpenAI) and Gemini were used solely for language editing of the manuscript. The authors take full responsibility for the final content.

## Supporting Information

FIG S1 Effects of exogenous dU on vegetative growth of *Pyricularia oryzae*.

(A) Representative colonies (top) and colony area (bottom) of *P. oryzae* grown on complete medium (CM) with the indicated concentrations of dU for 11 days. (B) Representative liquid cultures (top) and dry weight of mycelia (bottom) grown in liquid CM with the indicated concentrations of dU for 6 days. Data represent means ± SD from three independent biological replicates. ns, not significant; * *P* < 0.05 (one-way ANOVA followed by Dunnett’s multiple comparisons test).

FIG S2 Effect of dU treatment on appressorium diameter of *Pyricularia oryzae*.

(A) Workflow for appressorium detection and diameter measurement using the Hough Circle Transform plugin in Fiji. (B) Schematic cross-section of an appressorium attached to the plant surface, illustrating the basis for diameter measurement. Adapted from 33, Fig. 2. (C) Violin plots showing appressorial diameter at 24 h in the control, 2 µM dU, and 5 µM dU groups. Individual data points are overlaid. The thick horizontal line indicates the median, and the dashed lines indicate the interquartile range (IQR). Statistical significance was determined using the Kruskal-Wallis test with Dunn’s post-hoc test for multiple comparisons (* *P* < 0.05; ns, not significant), as the data did not pass the D’Agostino-Pearson normality test (*P* < 0.05).

FIG S3 Intracellular localization and buffer dependence of dU-metabolizing activity in strain C3.

(A) Total dU-metabolizing activity was compared between the crude intracellular extract and the culture supernatant of strain C3. (B) dU-metabolizing activity of the crude intracellular extract was examined in potassium phosphate (KPi) and Tris-HCl buffers at the indicated pH values.

FIG S4 Maximum-likelihood phylogenetic tree of DeoA-like proteins from Enterobacterales.

The tree was inferred from a trimmed amino acid alignment using IQ-TREE v3.0.1 under the Q.PFAM+I+R6 model. The DeoA sequence from *Enterobacter* sp. strain C3 is highlighted in blue. Taxon names are followed by protein accession numbers, and the sequences used in the analysis are listed in Table S2. The scale bar indicates amino acid substitutions per site.

FIG S5 Calibration of the gel filtration column using standard protein markers.

Retention times (RT) for the protein standards were as follows: bovine serum albumin (67 kDa), 9.39 min; ovalbumin (43 kDa), 10.07 min; chymotrypsinogen A (25 kDa), 12.59 min; and ribonuclease A (13.7 kDa), 13.70 min.

Table S1 Medium components of dUMM.

Table S2 DeoA-like proteins used for phylogenetic analysis.

Table S3 Primer used in this study.

